# Free energies of stalk formation in the lipidomics era

**DOI:** 10.1101/2021.06.02.446700

**Authors:** Chetan S. Poojari, Katharina C. Scherer, Jochen S. Hub

**Affiliations:** Theoretical Physics and Center for Biophysics, Saarland University, Saarbrücken, Germany

## Abstract

Many biological membranes are asymmetric and exhibit complex lipid composition, comprising hundreds of distinct chemical species. Identifying the biological function and advantage of this complexity is a central goal of membrane biology. Here, we study how membrane complexity controls the energetics of the first steps of membrane fusions, that is, the formation of a stalk. We first present a computationally efficient method for simulating thermodynamically reversible pathways of stalk formation at near-atomic resolution. The new method reveals that the inner leaflet of a typical plasma membrane is far more fusogenic than the outer leaflet, which is likely an adaptation to evolutionary pressure. To rationalize these findings by the distinct lipid compositions, we computed ~200 free energies of stalk formation in membranes with different lipid head groups, tail lengths, tail unsaturations, and sterol content. In summary, the simulations reveal a drastic influence of the lipid composition on stalk formation and a comprehensive fusogenicity map of many biologically relevant lipid classes.

## Introduction

Eukaryotic cellular membranes contain more than ten lipid classes, while each class comprises hundreds of different chemical species.^1,2^ The complexity of membranes is further increased by the membrane asymmetry, that is, by distinct lipid compositions in the two leaflets. In the plasma membrane of mammals, sphingolipids are typically enriched in the outer leaflet of the plasma membrane, whereas phosphatidylethanolamine (PE) and phosphatidylserine (PS) are enriched in the inner leaflet.^3^ Apart from different head groups, lipid species differ by the length and unsaturation of the fatty acid tails. Recent lipidomics studies showed that polyunsaturated lipids are strongly enriched in the inner leaflet.^4^ Understanding why biological cells synthesize and maintain this complex lipid repertoire, that is, defining the biological function and advantage of specific lipid compositions, remains a central goal of membrane biophysics.

Here, we study how complex lipid compositions control the early stages of membrane fusion. Fusion is critical for many processes that involve the transport of cargos across membranes such as exocytosis, neurotransmission, infection of cells by enveloped viruses, fertilization, and intracellular transport.^5–8^ Membrane fusion occurs with the help of fusion proteins because fusion requires the two opposing lipid membranes to overcome a large dehydration barrier. The process starts with two membranes in the lamellar phase at the equilibrium distance of 2-3 nm, followed by bridging of two opposing membranes by lipid acyl chains to establish a point-like protrusion. The lipids of the two contacting, proximal leaflets mix to form the initial transient hemifusion stalk. The stalk expands to allow the formation and expansion of a fusion pore.^9–11^

Because intermediate structures along the fusion pathway involve highly curved membranes, the intrinsic curvature of lipids influences the kinetics of fusion. For instance, when added to the proximal leaflets, inverted cone-shaped lysophosphatidylcholine (LPC) inhibits stalk formation because it is incompatible with the large negative curvature at the stalk rim. Inversely, cone-shaped unsaturated PE lipids or diacylglycerol promote stalk formation.^10,12,13^ Here, the unsaturated fatty acids render the lipids more cone-shaped because the double bonds increase the tail disorder and, hence, the effective volume of the lipid tails. ^13^ However, in addition to such effect by the geometric lipid shape, the increased abundance of double bonds increases the conformational flexibility of the lipids, thereby allowing the membrane to adapt more easily to curvature, ^14^ which may further promote fusion.

Theoretical and computational approaches have established possible fusion pathways and the underlying free energy landscapes, however, only few studies focused on the role of the lipid composition during fusion. Early studies employed continuum descriptions, which model the role of lipids in terms of the effective spontaneous curvature and membrane rigidity, ^15^ as well as minimal coarse-grained lipids models in conjunction with Monte-Carlo, field-theoretic, or Brownian dynamics methods.^16–18^ Complementary, to account for the chemical specificity of lipid–lipid interactions, fusion was studied with atomistic and coarse-grained (CG) molecular dynamics (MD) simulations,^19–21^ discussed in several excellent reviews.^22–24^ PE lipids were found to enhance fusion during simulations as expected from their negative intrinsic curvature,^25–27^ however the effects of other head groups, sterols, or degrees of unsaturation have hardly been considered despite their abundance in biological membranes.

Computationally efficient free energy calculations of fusion require the definition of one or several reaction coordinates (or order parameters) along the fusion pathway; however, finding good coordinates for complex transitions is non-trivial. With poor coordinates, hysteresis problems emerge, barriers may be integrated out, and simulations proceed along non-reversible pathways. To avoid such problems, a recent elegant MD study used the string method to optimize the minimum free energy pathway of stalk formation, parametrized by the three-dimensional density of the apolar lipid tail beads.^27^ However, because the string method is computationally demanding, it becomes prohibitive for high-throughput studies. In this study, we introduce a reaction coordinate for stalk formation that allows computationally highly efficient and thermodynamically reversible free energy calculations of stalk formation using common umbrella sampling simulations. The potentials of mean force (PMFs) along the coordinate, computed with Martini coarse-grained models, show that the inner leaflet of a common plasma membrane is far more fusogenic than the outer leaflet. To rationalize these finding, we screen the fusogenicity of lipids by varying the hydration level between the two membranes, the lipid headgroup size and charge, acyl chain length, and unsaturation levels. In addition to phosphatidylcholine (PC), PE, PS, and phosphatidylglycerol (PG) lipids, we also considered cholesterol, lysolipids, fatty acids in protonated and deprotonated forms, phosphatic acid, ceramide, diacylglycerol, and sphingomyelin. The calculations provide a comprehensive view on the role of lipid complexity during the first steps of membrane fusion.

## Results

### Highly efficient free energy calculations of stalk formation

Potential of mean force (PMF) calculations may provide the free energy along reaction path in a computationally efficient manner. However, they require the choice of a good reaction coordinate (or order parameter); otherwise, undesired hysteresis effects may emerge, or energy barriers may be integrated out. We propose using a reaction coordinate for stalk formation that was originally introduced to study the formation of aqueous pores over membranes.^28,29^ The coordinate, named “chain coordinate” *ξ*_ch_, quantifies the degree of connectivity between two compartments. In brief, *ξ*_ch_ is defined with the help of cylinder that spans the head group and water regions of the proximal compartment (Fig. S1). The cylinder is decomposed into *N_s_* slices, and *ξ*_ch_ is defined as the fraction of slices that are filled by apolar lipid tail beads. By pulling the system along the coordinate, the slices are filled one-by-one, thereby forming a continuous apolar connection between the two fusing membranes, as required for stalk formation (Fig. 1A–C). More details are provided in the supporting material.

**Figure 1.**
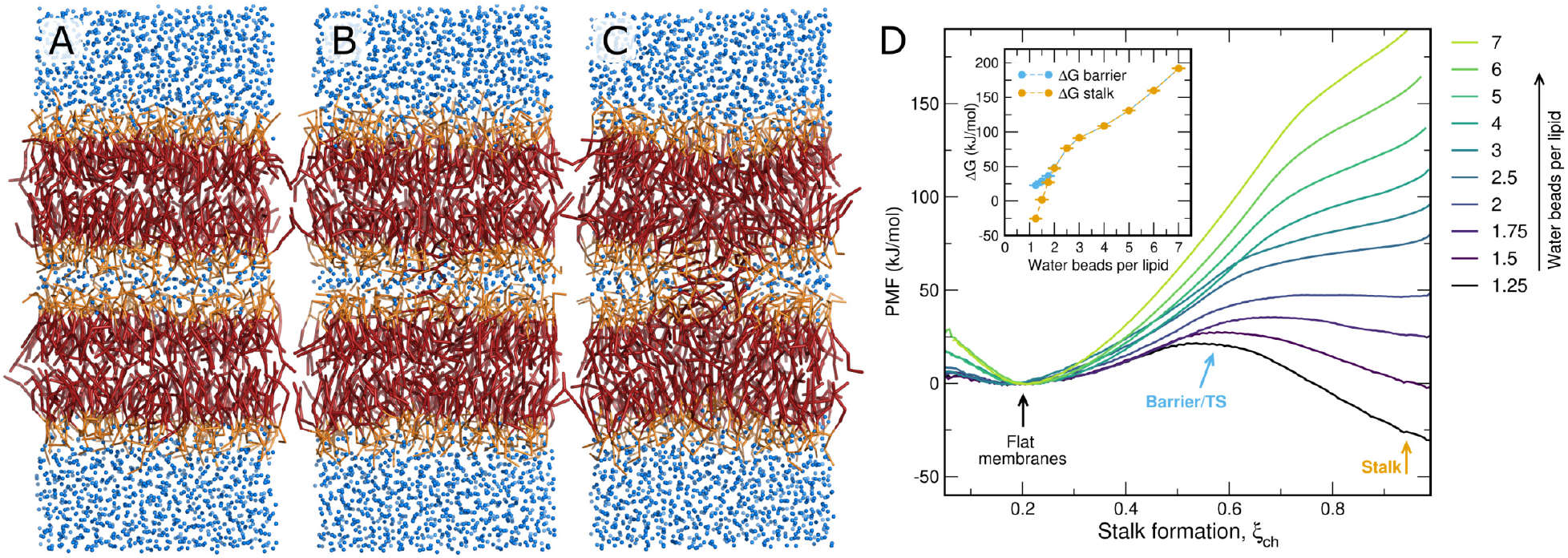
(A–C) Martini simulation system with membranes of pure POPC and 1.5 water beads per lipid in the proximal water compartment, showing representative frames of the (A) the flat membrane (B) the transition state (TS), and (C) the stalk. (D) PMFs of stalk formation for membranes of pure POPC, with increasing amount of water in the proximal compartment (see color code). Inset: Free energy of the stalk Δ*G*_stalk_ and of the stalk nucleation barrier (Δ*G*_barrier_, if present) versus water beads per lipid in the proximal compartment, as taken from the PMFs.

We used umbrella sampling along *ξ*_ch_ to compute the PMF of stalk formation in conjunction with the coarse-grained Martini lipid force field. Figure 1D presents PMFs of stalk formation across membranes of POPC lipids using different degrees of hydration in the proximal water compartment, spanning 1.25 to 7 water beads per lipid, corresponding to 5 to 28 water molecules per lipid according to the 4:1 mapping of Martini. Here, the length of the *ξ*_ch_-defining cylinder was chosen such that *ξ*_ch_ ≈ 0.2 corresponds to flat unperturbed membranes, implying that 20% of the cylinder slices are filled by lipid tail beads. *ξ*_ch_ ≈ 1 corresponds to a fully formed stalk. Evidently, free energy of the stalk Δ*G*_stalk_ relative to the flat membranes greatly depends on the degree of hydration, spanning values from −30kJ/mol up to 180kJ/mol (Fig. 1D, inset). With fewer than 2 water beads per lipid, a barrier or transition state emerges, indicating that the stalk is metastable (long-living); with less than 1.5 water beads per lipid the stalk even forms the free energy minimum. The marked dependence of Δ*G*_stalk_ on the degree of hydration agrees with a recent simulation study, which used the string method together with the three-dimensional (3D) lipid tail density as order parameter to derive the minimum free energy path of stalk formation.^27^ However, with the identical simulation system, the free energies for the stalk suggested by our PMFs are significantly lower (Fig. S2), possibly because our stalk state given by *ξ*_ch_ ≈ 1 includes more conformational freedom than a stalk definition via the 3D density used in Ref. 27.

The Martini model allows semi-quantitative simulations, suggesting that Martini yields *trends* often correctly;^30^ however, the exact free energy values must often be taken with care and may depend on the Martini version. To test how the stalk free energies depend on the Martini version, we re-computed the PMFs of stalk formation for POPC with the beta-3.2 release of Martini 3.0 (3.0beta) instead of Martini 2.2 (Fig. S3). Evidently, the shape of the PMFs and the dependence on hydration are well preserved among different Martini versions. However, the free energy of the stalk with Martini 3.0beta is reduced by ~30 kJ mol^-1^ relative to Martini 2.2, suggesting that the newer Martini version is more fusogenic. Simulations with four additional lipid types confirm this trend (Fig. S4). This finding suggests that the free energies reported in this study should be interpreted in terms of trends (with hydration, degree of unsaturation etc.) and not in terms of precise free energy values.

### Kinetics of stalk formation agree with the PMFs

As a critical test for the quality of the reaction coordinate and, thereby, for the validity of the PMFs, we carried out free simulations of stalk formation and stalk closure (Fig. 2). We simulated a system for which Δ*G*_stalk_ ≈ 0 according to the PMF (Fig. 2B). Among a total simulation time of 800 μs, we observed 8 transitions of stalk opening and 7 transitions of stalk closure, corresponding to rates of *k*_stalk_ = 16 ms^-1^ and *k*_closure_ = 23 ms^-1^, respectively. Hence, the free simulations suggest a free energy of stalk formation of Δ*G*_stalk_ = -*k_B_T*ln(*k*_stalk_/*k*_closure_) = 0.9 kJ/mol. The excellent agreement with the PMF suggests that the PMF is not affected by hysteresis problems but reports the correct Δ*G*_staik_ as given by the Martini force field. Using transition state theory, the rates are expected to follow *k* = *v*e^-Δ*G*‡/*k_B_T*^, where *v* is the attempt frequency, Δ*G*^‡^ the free energy barrier in Fig. 2B, and *k_B_* and *T* are the Boltzmann constant and temperature, respectively. The simulations yield *v* ≈ 0.3ns^-1^, i.e., one attempt per 3ns, which corresponds approximately to the time scale that lipids take for traveling a lipid–lipid distance. It is interesting to note that this natural time scale for lateral lipid diffusion appears also as time scale for attempts of stalk formation.

**Figure 2:**
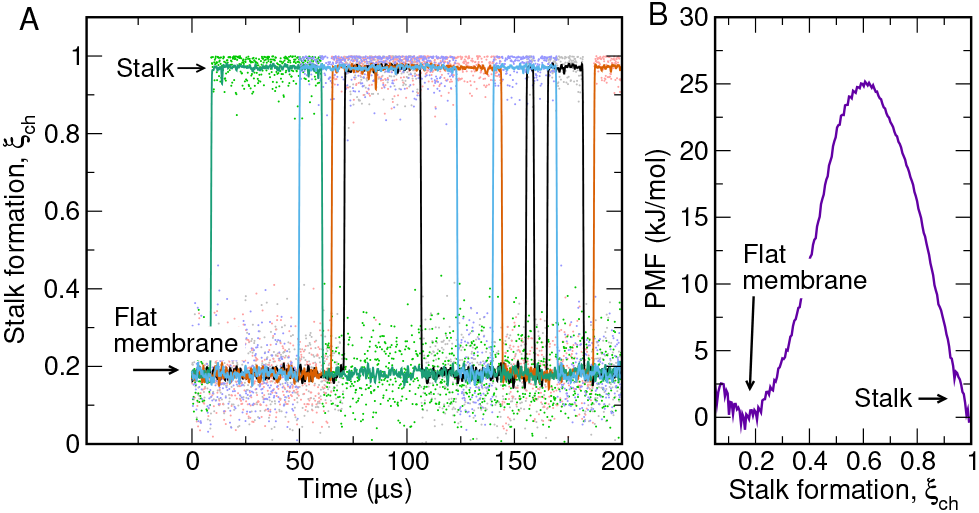
(A) Free simulations of stalk formation and closure. Colors indicate four independent simulations of 200 μs each. *ξ*_ch_ ≈ 0.2 and *ξ*_ch_ ≈ 1 correspond to the flat membrane and the stalk, respectively. Evidently, 8 transitions of stalk formation and 7 transitions of stalk closure occurred within a total simulation time of 800 μs. The simulation system contained purely POPC lipids and 1.8 water beads per lipid between the proximal leaflets. Lipids were modelled with the beta-3.2 release of Martini 3.0. (B) PMF of stalk formation for the same system, revealing a free energy of stalk formation of Δ*G*_stalk_ ≈ 0 kJ/mol, in excellent agreement with the free simulations.

### The inner leaflet of the plasma membrane is far more fusogenic than the outer leaflet

Many critical fusion events occur with the plasma membrane. During exocytosis, transport vesicles fuse with the plasma membrane, where they first form a stalk with the *inner* plasma membrane leaflet. During viral infection, the envelope of certain viruses fuses with the plasma membrane, thereby forming a stalk with the *outer* leaflet. Like many biological membranes, the plasma membrane is asymmetric, i.e., the two leaflets exhibit different lipid compositions. Further, the plasma membrane reveals a complex lipid composition, including various head group types, tail lengths, degrees of tail unsaturation, and steroid content.^4^

To reveal how the lipid composition of a complex biological membrane determines the free energies of stalk formation, we set up systems with symmetric membranes, but with the lipid composition mimicking either the outer or inner leaflet of a mammalian plasma membrane.^4^ The membranes contained phosphatidyl-choline (PC), -ethanolamine (PE), – inositol (PI), -serine (PS), sphingomyelin (SM), and cholesterol, as well a various tail lengths and degrees of unsaturation, taken from a recent lipidomics study^4^ (Fig. 3D/E, Tables 1 and S1). The PMFs of stalk formation were again computed for various degrees of hydration in the proximal compartment quantified by the number of water beads per membrane area (Figs. 3A/B and S5). Remarkably, the free energy of stalk formation between membranes with the outer leaflet composition is larger by ~50kJ/mol as compared to the membrane with the inner leaflet composition, irrespective of the degree of hydration (Figs. 3C, yellow dots). The same trend is observed for the free energy barrier for stalk formation (if present, Figs. 3C, blue dots). Hence, the inner leaflet is far more fusogenic than the outer leaflet of the plasma membrane. These different fusogenicities may reflect adaptation to the evolutionary pressure: Efficient fusion with the inner leaflet is required for exocytosis, in particular for rapid fusion of synaptic vesicles with the plasma membrane of the synapse. In contrast, an increased resistance against fusion with the outer leaflet may protect the cell against viral infection.

**Figure 3:**
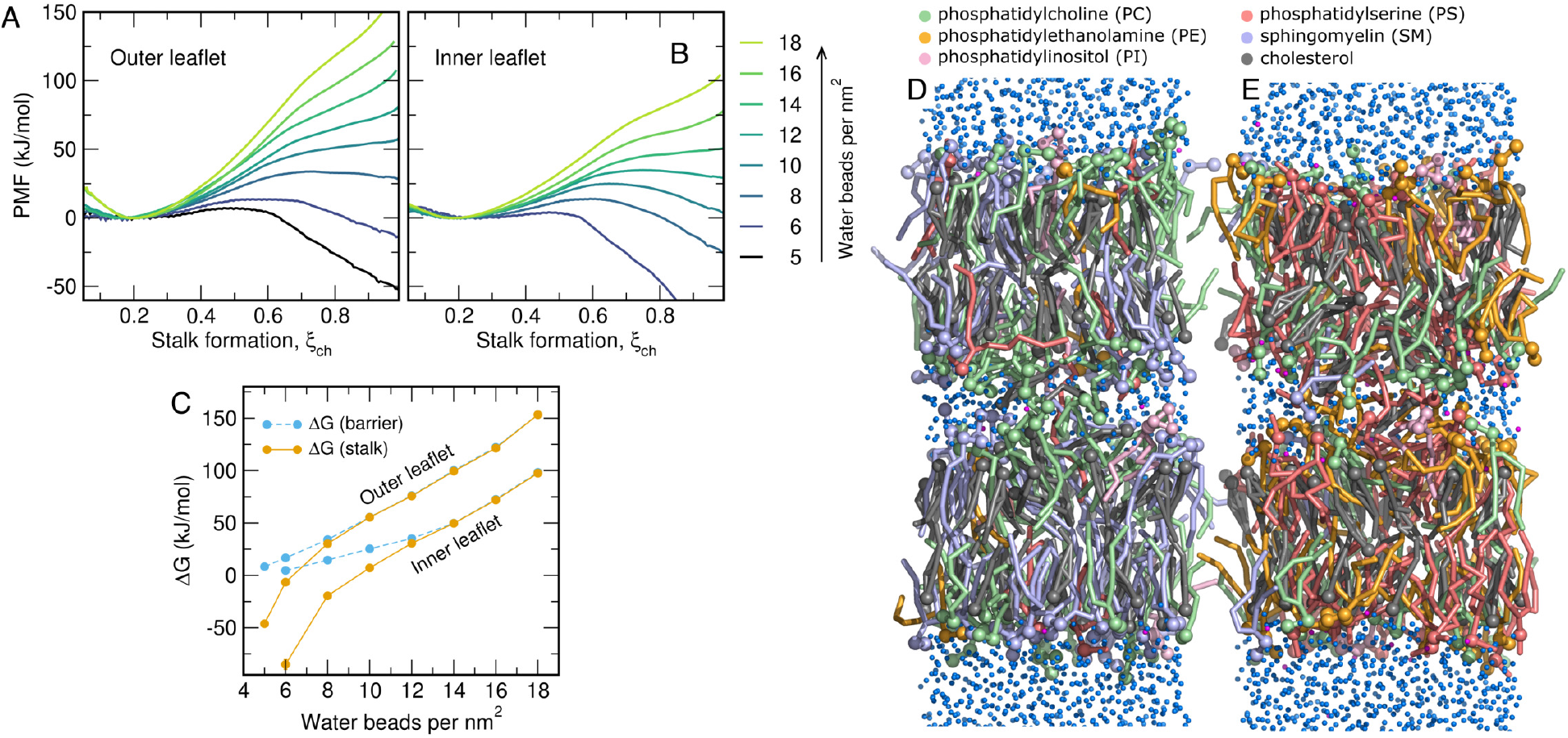
(A/B) PMFs of stalk formation with lipid composition taken from the plasma membrane (A) outer leaflet or (B) inner leaflet. *ξ*_ch_ ≈ 0.2 and *ξ*_ch_ ≈ 1 correspond to the states with planar membranes and with the stalk, respectively. PMFs were computed with increasing hydration between the proximal leaflets, between 5 and 18 water beads per nm^2^. (C) Free energy of the stalk (orange) and of the barrier for stalk formation (blue), taken from the PMFs in A/B. (D) Simulation frame of the open stalk in simulation with outer leaflet composition and (E) inner leaflet composition. Lipids are shown as colored sticks, headgroup beads as spheres (for color code, see legend). Water and sodium beads are shown as small blue and magenta spheres, respectively.

**Table 1:** Summary of lipid composition and properties in models for the outer and inner leaflet.

|  | Outer | Inner |
| --- | --- | --- |
| PC | 25% | 15% |
| PE | 2% | 13% |
| SM | 25% | 2% |
| PI | 2% | 2% |
| PS | 3% | 25% |
| Cholesterol | 43% | 43% |
| Saturated beads | 89% | 77% |
| Unsaturated beads | 11% | 23% |
| Av. tail length (beads) | 3.9 | 4.2 |

### Free energies of stalk formation are strongly influenced by tail unsaturation, tail length, and headgroup type

Why is the inner plasma membrane leaflet more fusogenic than the outer leaflet? As listed in Tables 1 and S1, the inner leaflet model contains more PE and PS lipids, whereas the outer leaflet contains more PC and SM lipids. In addition, the lipid tails of the inner leaflet are more unsaturated as compared to the outer leaflet, as given by the fraction of beads modeling unsaturated tails (23% versus 11%), and the lipid tails of the inner leaflet are longer on average (4.2 versus 3.9 beads). Owing to this complexity, it is difficult to extract the key lipid properties underlying the distinct fusogenicities.

To disentangle the influence of tail unsaturation, tail length, and type of head group we computed 110 additional PMFs of stalk formation. These calculations were accessible only thanks to the computational efficiency of the PMF calculations. We systematically varied either (i) the degree of unsaturation by simulating fully saturated up to poly-unsaturated tails, (ii) the tail length from 3 to 6 coarse-grained beads, thereby modeling approximately 14 to 26 carbon atoms per tail, or (iii) the type of head group between PS, PG, PC, and PE (Fig. 4A–C). The PMFs were again computed for various degrees of hydration between 4 and 18 water beads/nm^2^ (Figs. S6 and S7). Molecular representations of the stalks are shown in Figs. S8, S9, and S10.

**Figure 4:**
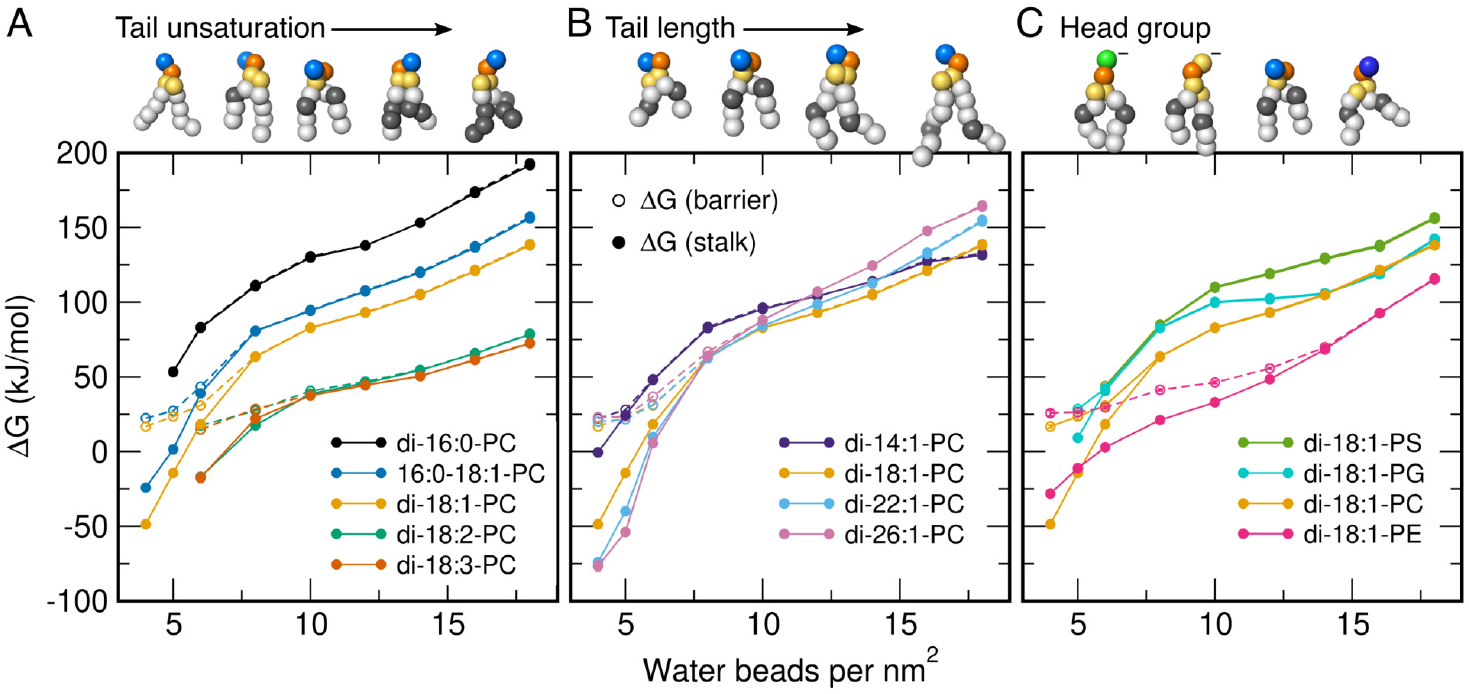
Free energies of stalk formation Δ*G*_stalk_ (solid circles) for single-lipid membranes composed of various lipid types, plotted as a function of increasing hydration between the proximal leaflets. Legends indicate the approximate atomistic correspondence of the Martini models. Open circles show the free energy barrier for stalk formation (if present). Beads of Martini lipid models are colored as follows: hydrophobic saturated (white), hydrophobic unsaturated (grey), glycerol (yellow), phosphate (orange), choline (light blue), serine (green), ethanolamine (dark blue).

Evidently, apart from the degree of hydration, also the lipid type strongly influences the free energy of the stalk Δ*G*_stalk_ and the free energy barrier for stalk formation. Specifically, saturated tails as found in DPPC (di-16:0-PC) disfavor stalk formation, whereas polyunsaturated tails in lipids such as di-18:2-PC greatly favor stalk formation (Fig. 4A). In addition, increasing the tail length may stabilize the stalk at low hydration of ≤6 water beads/nm^2^ for which the stalk is energetically accessible (Fig. 4B). At increased hydration, however, for which the stalk is unstable, longer tails further destabilize the stalk. Likewise, the type of head group may influence Δ*G*_staik_: the anionic head groups of dioleoyl-PS (DOPS) and DOPG disfavor stalk formation by ~25kJ/mol relative to the zwitterionic head group of DOPC (Fig. 4C). These findings cannot be explained merely the geometric shape of the lipids because PS headgroups are smaller than PC headgroups, whereas PG headgroups are larger than PC headgroups. Hence, the electrostatic repulsion between the anionic lipids plays a distinct role in destabilizing the stalk, possibly by effectively inducing a more positive spontaneous membrane curvature. The small zwitterionic PE headgroup facilitates stalk formation relative to the larger PC headgroup by up to 50 kJ/mol, as one may expect from the cone shape of PE lipids that may stabilize the strong negative curvature in the stalk. This trend is only inverted at very low hydration of ≤5 water beads/nm^2^, where stalk formation between PC membranes is more favorable than between PE membranes (Fig. 4C, yellow and magenta dots). Simulations with PE and PC membranes with alternative tails confirmed that, at most degrees of hydration, PE headgroups typically facilitate stalk formation relative to PC (Fig. S6).

Taken together, these data demonstrate that the free energy cost for stalk formation is controlled by a range of lipid properties. Namely, stalk formation is facilitated by (i) increased tail unsaturation, (ii) by longer tails at low hydration between the fusing membranes, (iii) by zwitterionic relative to anionic lipids, and (iv) by the small PE instead of the larger PC headgroup (at most degrees of hydration).

### Δ*G*_staik_ scales linearly with lipid concentration in lipid mixtures

Next, we investigated how the lipid type and concentration influence stalk formation in lipid mixtures. To this end, we set up addition 76 simulation systems containing POPC as reference lipid plus one type of lipid with increasing mole fraction. A constant degree of hydration with 6 water beads/nm^2^ was used. In addition to the PC, PG, PS, and PE lipids considered above, and to obtain a comprehensive view on the influence of lipids on stalk free energies, we considered a wide range of additional lipids: cholesterol, lyso lipids, fatty acids, phosphatidic acid (PA), ceramide, diacylglycerol, and sphingomyelin (SM).

Figure 5 presents Δ*G*_stalk_ as function of mole fraction *x*_lipid_ for various lipid types, where *x*_lipid_ = 0% corresponds to pure POPC. Typical PMFs are presented in Fig. S11. Evidently, mixing a second type of lipid into a POPC membrane may greatly stabilize or destabilize the stalk, while Δ*G*_stalk_ depends nearly linearly on the mole fraction of the second lipid. Here, the slope of the Δ*G*_stalk_-*x*_lipid_ curves quantify the sensitivity of stalk formation upon the addition of a second lipid.

**Figure 5:**
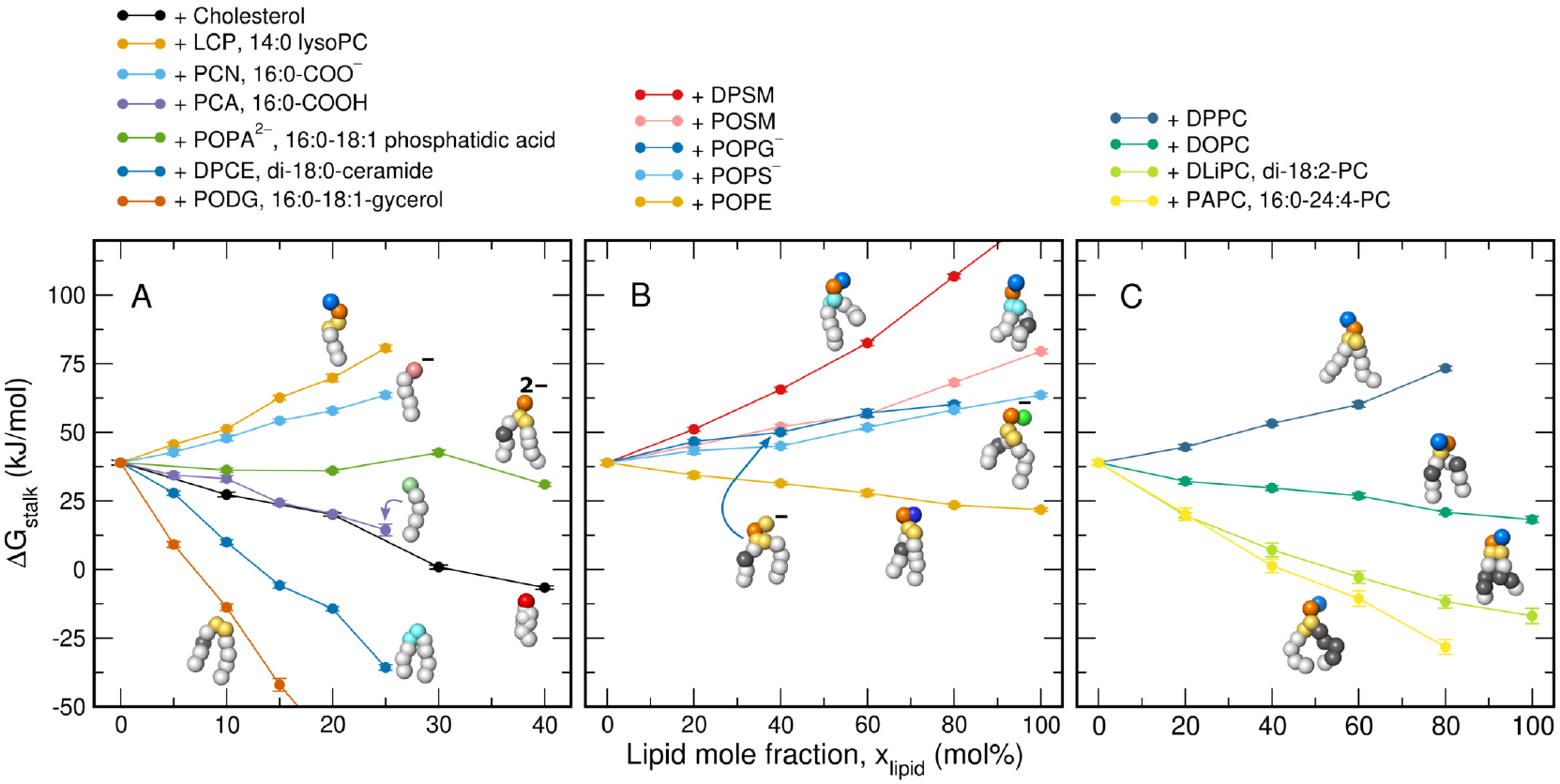
Free energies of stalk formation Δ*G*_stalk_ for binary lipid mixtures of POPC plus one additional type of lipid (for color code, see legend). 0mol% correspond to pure POPC. Martini beads are colored as with the scheme used for Fig. 4 and as follows: hydroxyl (red), carboxylate (pale red), carboxyl (pale green), sphingosine (cyan).

As expected, the geometric shape of the lipid, that is whether the lipid is cone- or inverted cone-shaped, affects Δ*G*_stalk_. Only small amounts of ceramide and diacylglycerol greatly stabilize stalks, compatible with their highly negative intrinsic curvature (Fig. 5A, dark blue, dark orange). On the contrary, single-chained lysoPC greatly destabilizes the stalk, compatible with its large positive curvature (Fig. 5A, orange). Single-chained fatty acids stabilize the stalk in the protonated form 16:0-COOH, however they destabilize the stalk in the deprotonated form 16:0-COO^-^ (Fig. 5A, light blue, purple). Owing to the strongly increased pK_a_ of fatty acids in lipid membranes relative to bulk water, the protonated, stalk-stabilizing form is predominantly present in biological membranes.^31^

In agreement with the findings for single-component membranes presented above, Δ*G*_stalk_ is modulated by replacing PC head groups of POPC with other head groups while maintaining the palmitoyl-oleoyl tails. Whereas replacing PC with PA has only a small effect on Δ*G*_stalk_, replacing PC with PE favors stalk formation (Fig. 5A, green, Fig. 5B, orange). In contrast, replacing PC with the anionic PS or PG head groups disfavors stalk formation (Fig. 5B, light and dark blue). Likewise, sphingomyelin disfavors stalk formation, in particular the fully saturated DPSM. These findings are rationalized by the tendency of sphingomyelin of increasing the order of the aliphatic tails, thereby disfavoring the formation of a defect such as a stalk. In addition, the Δ*G*_stalk_ values from lipid mixtures confirm that the addition of (poly-)unsaturated lipid tails facilitate stalk formation, whereas the addition of saturated tails hinder stalk formation (Fig. 5C). Finally, the addition of cholesterol strongly stabilizes the stalk, by up to 45 kJ/mol at 40 mol% cholesterol, in qualitative agreement with results from X-ray diffraction. ^32^

### Stalk stabilization correlates with specific lipid enrichment in the stalk, except for cholesterol

To shed light on the molecular mechanism by which different lipids influence Δ*G*_stalk_, we computed the lipid densities in simulations with a fully formed stalk (Figs 6A/B and S12– S15). The overall shape of the stalk is largely independent of the lipid composition, except that energetically unstable stalks are slightly thinner as compared to stalks that form the free energy minimum. However, the lipid composition strongly influences the spatial distribution of the individual lipid types. For instance, the stalk-stabilizing DLiPC is enriched at the stalk, whereas stalk-destabilizing DPSM is depleted at the stalk (Figs 6A/B, left column). To test whether these trends hold for other lipids, we quantified the enrichment of a lipid in the stalk as *ρ*_stalk_/*ρ*_mem_ — 1, where *ρ*_stalk_ and *ρ*_mem_ denote the average lipid mass densities at the stalk and in a region of the flat membrane, respectively (Figs 6A, white and yellow boxes). Hence, a positive and negative values indicate lipid enrichment or depletion at the stalk, respectively. In addition, we quantified the influence of specific lipids on stalk formation as the slope of a linear least-square fit to the data in Fig. 5, denoted ΔΔ*G*_stalk_/*x*_lipid_.

**Figure 6:**
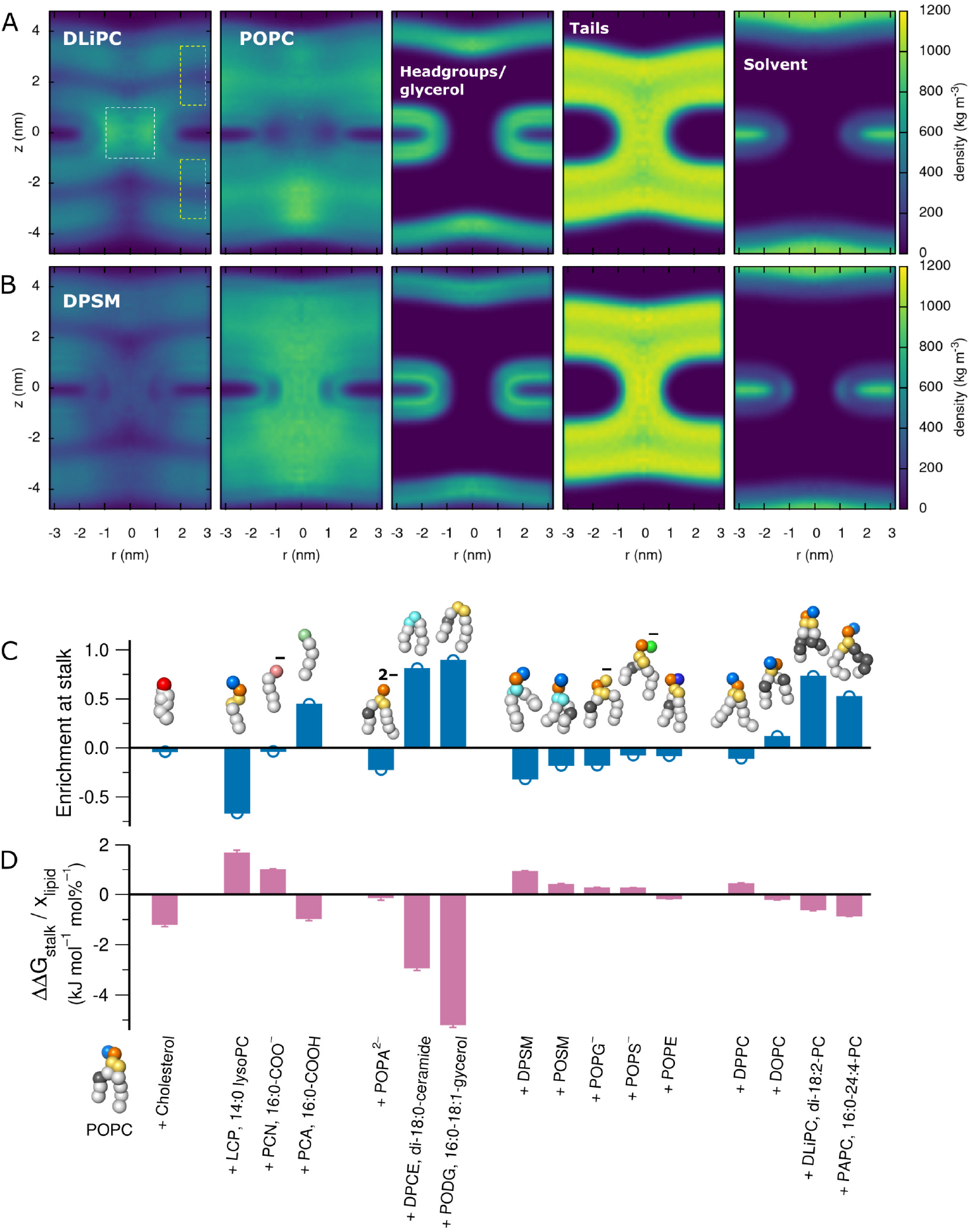
(A) Mass densities of (from left to right) DLiPC, POPC, headgroups plus glycerol beads, lipid tails, and solvent in the fully formed stalk in a system with a POPC:DLiPC ratio of 60:40. (B) Same as in panel (A), but for the DPSM:POPC 60:40 system. (C) Enrichment of lipids inside the stalk relative to the flat membrane: *ρ*_stalk_/*ρ*_mem_ — 1, where *ρ*_stalk_ and *ρ*_mem_ are the average densities in the white and yellow boxes in panel (A), respectively. Beads of Martini lipid models are colored as follows: hydrophobic saturated (white), hydrophobic unsaturated (grey), hydroxyl (red), glycerol (yellow), phosphate (orange), choline (light blue), carboxylate (pale red), carboxyl (pale green), sphingosine (cyan), serine (green), ethanolamine (dark blue). (D) Change of stalk free energy ΔΔ*G*_stalk_ per lipid concentration *c*_lipid_ upon addition of lipid to a POPC membrane.

Evidently, among nearly all lipids, lipid enrichment in the stalk anti-correlates with the ΔΔ*G*_stalk_/*x*_lipid_. Hence, lipids that either favor negative curvature, increased tail disorder, or both, may accumulate in the stalk, thereby stabilizing the stalk structure. However, there are notable outliers: cholesterol stabilizes the stalk without being enriched in the stalk. Instead, in line with previous reports,^33^ cholesterol is even slightly depleted at the stalk, possibly because the planar cholesterol molecule disfavors the increased tail disorder in the stalk. These findings suggest that cholesterol does not favor stalk formation owing to an intrinsic negative curvature of cholesterol, but more likely owing to its influence on the dehydration repulsion between the membranes.

Another outlier is diacyl-glycerol PODG, the by far most stalk-stabilizing lipid considered in this study. The lipid densities show that PODG accumulated not only inside the stalk, but mostly on top and below the stalk within the membrane core (Fig. S13). The accumulation within the membrane core is possible because the headgroup of diacyl-glycerol is only weakly polar. We hypothesize that similar effects play a role during biogenesis of lipid droplets, which contain large amounts of triacylglycerol and which are released from the ER membrane via a stalk-like intermediate.^34^

## Discussion

The marked difference between the outer and inner leaflet prompted us to screen free energies of stalk formation for a wide range of single-lipid membranes as well as for membranes of binary lipid mixtures. The results provided a comprehensive and quantitative fusogenicity map of lipids, as summarized in Fig. 6D. The data suggest that free energies of stalk formation are not determined by a single lipid property, but instead by a combination of molecular mechanisms: (i) Stalk formation is facilitated by increased tail disorder, as present in polyunsaturated lipids, whereas increased tail order hinders stalk formation, as common for sphingomyelin. (ii) As expected from previous studies,^35^ lipids with negative intrinsic curvature promote stalk formation as they may stabilize the large negative curvature of the stalk. Such mechanism applies to protonated fatty acids, to PE lipids and ceramide owing to their small head groups, as well as to polyunsaturated lipids owing to the increased tail volume. A counter example is given by lysoPC that hinders stalk formation owing to its positive intrinsic curvature. ^35^ (iii) Likewise, we find that anionic headgroups counteract stalk formation. The simulations demonstrate that anionic headgroups counteract stalk formation, as implied by increased Δ*G*_stalk_ for PS or deprotonated fatty acids despite of their small head groups. In line with this observation, POPA hardly affects Δ*G*_stalk_ despite the very small headgroup. These trends are likewise rationalized by the electrostatic repulsion between headgroups, which effectively disfavor negative curvature.^36^ However, since polar interactions are less well represented by the Martini coarse-grained force field as compared to apolar interactions, it will be interesting to test these predictions by experiments and by atomistic simulations. (iv) Cholesterol facilitates stalk formation. However, since cholesterol is not enriched in the stalk, the stalk-stabilizing effect of cholesterol can not rely on a putative stabilization of the negative curvature in the stalk, as suggested previously. ^37^ The effect of cholesterol is also surprising considering that cholesterol increases the tail order of membranes, similar to the sphingomyelin that, in contrast to cholesterol, increases Δ*G*_stalk_. Therefore, we hypothesize that cholesterol may promote stalk formation by reducing the hydration repulsion,^38^ thereby facilitating the formation of the first contacts between the fusing leaflets.

By simulating stalk formation between membranes with complex lipid composition, we found that the inner leaflet of a typical mammalian plasma membrane^4^ is more fusogenic than the outer leaflet by ~50kJ/mol. These properties may be an adaption to evolutionary pressure: a fusogenic inner leaflet facilitates exocytosis, thereby increasing the fusion rates and reducing the required energy consumption by fusion proteins. A less fusogenic outer leaflet might hinder infection by enveloped viruses that fuse with the plasma membrane. The fusogenicity of lipids in Fig. 6D provides the molecular rationale for the distinct fusogenicities of the inner and outer leaflets. Namely, the increased fusogenicity of the inner leaflet is mainly caused by the increased number of polyunsaturated tails, mainly present as PS and PE lipids, and, to a lower degree, by the large PE content (cf. Table 1). The outer leaflet counteracts fusion mainly owing to the large sphingomyelin content, fewer polyunsaturated tails, and increased PC content relative to PE.

These conclusions were obtained with novel reaction coordinate for stalk formation, which allowed computationally highly efficient free energy calculations of stalk formation along thermodynamically reversible pathways. The PMFs converge rapidly, are not affected by hysteresis problems, and they yield the true free energy difference between the lamellar and the stalk state for the given force field, as confirmed by free simulations. One PMF required less than 7 hours on an inexpensive server equipped with a 6-core CPU and a consumer graphics card, hence allowing for high-throughput calculations. Our PMF calculations complement computationally more elaborate methods such as the string method, which is capable of finding the minimum free energy path of stalk formation without the need of identifying a good reaction coordinate. ^27^ The computational efficiency further relied on the use of the Martini coarse-grained force field that achieves a speedup in sampling by a factor of ~1000 relative to atomistic simulations due to fewer particles, longer integration time step, and accelerated lipid diffusion. However, with the new reaction coordinate, free energy calculations of stalk formation with atomistic MD simulations are now within reach. This will allow us to test whether coarse-grained simulation primarily provide trends or whether they also provide quantitatively precise predictions of free energies during fusion.

While this study focused on stalk formation, it is evident that other stages of the fusion process are influenced by the lipid composition as well. Stalk widening as well as opening and expansion of the fusion pore involve highly curved membranes and are therefore controlled by lipid curvature.^10,39^ Hence, future studies should quantify the role of complex lipid compositions on such later stages of fusion, in addition to lipid effects on the dehydration repulsion, which may add a free energy offset to the free energies of stalk formation computed here.^27^ Further, it will be interesting to test whether lipid–protein interactions guide a local enrichment of fusogenic lipids and, thereby, further facilitate fusion.

## Conclusions

Using a newly developed method for computing free energies of stalk formation Δ*G*_stalk_, we presented a comprehensive and quantitative fusogenicity map of lipids. The simulations showed that the lipid composition of membranes may modulate Δ*G*_stalk_ by 100kJ/mol or even more. A range of lipid properties facilitate stalk formation, including increased tail unsaturation, longer tails, smaller head groups, and cholesterol content. In contrast, stalk formation is hindered by anionic lipids and by lipids that increase the tail order such as sphingomyelin. We found that the lipid composition of the inner leaflet of a mammalian plasma membrane greatly favors stalk formation relative to the outer leaflet. The distinct fusogenicities of the two leaflets is likely an adaptation to physiological requirements and is mainly rationalized by the different content of polyunsaturated lipids, sphingomyelin, and PE lipids.

## Methods Summary

MD simulations were carried out with Gromacs, version 20 20.3,^40^ and with an in-house modification of Gromacs 2018.8 that implements the harmonic restraint along the reaction coordinate *ξ*_ch_, originally introduced to study pore formation over membranes.^28,29^ PMFs were computed with umbrella sampling using 19 umbrella windows and simulating each window for 200 ns. Interactions were described with the Martini coarse-grained force field version 2.2 if not stated otherwise.^41^ Details on the reaction coordinate, simulation setup, parameters, and analysis are provided in the supporting material.

## Acknowledgement

We thank Yuliya Smirnova and Jelger Risselada for insightful discussions, for sharing the PChd200 and PCh220 simulation systems, and for critical comments on the manuscript. This study was supported by the Deutsche Forschungsgemeinschaft via SFB 1027/B7.

