## Supplementary material for "Free energies of stalk formation in the lipidomics era": Methods and Supporting Figures S1-S16

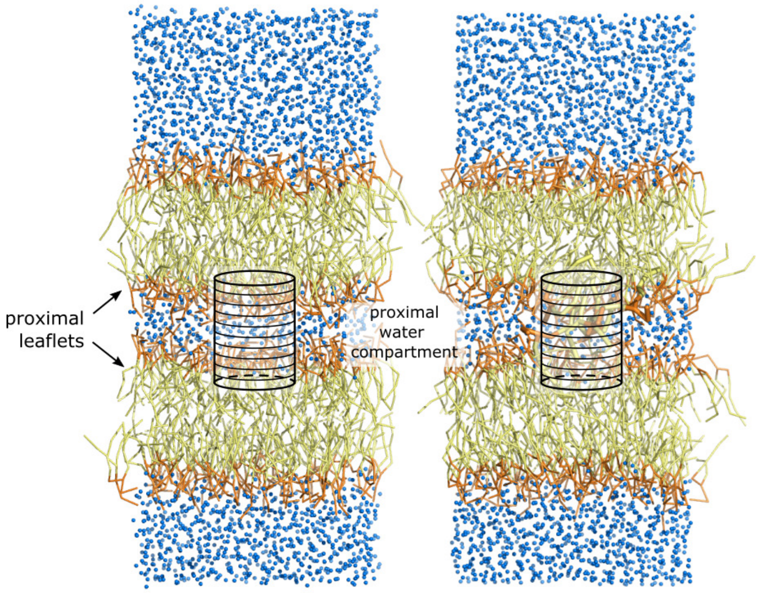


Figure S1: Illustration of a double-membrane system composed of POPC and the cylinder used to define the "chain coordinate" $\xi_{\mathrm{ch}}$. The coordinate $\xi_{\mathrm{ch}}$ is defined as the fraction of cylinder slices that are filled by apolar lipid tail beads. Tail beads are shown as yellow sticks, headgroup and glycerol beads as orange sticks, and water as blue spheres. Left: flat membrane, where only few cylinder slices is filled by tail beads ($\xi_{\mathrm{ch}} \approx 0.2$). Right: stalk, where all cylinder slices are filled by tail beads ($\xi_{\mathrm{ch}} \approx 1$). Lipids contributing to the stalk are highlighted by thicker sticks.

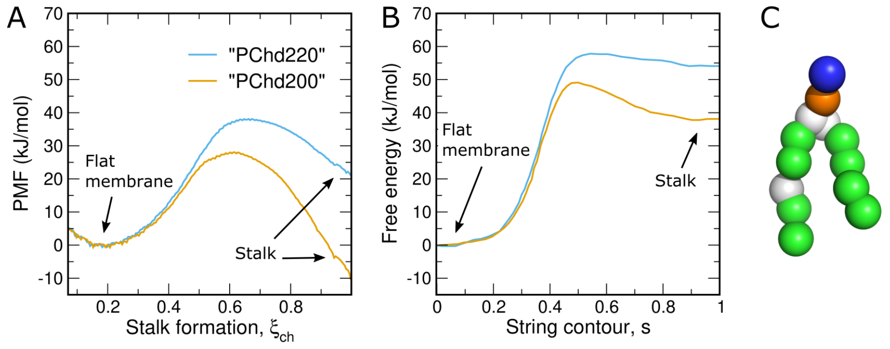


Figure S2: Comparison of (A) the PMF along the chain coordinate $\xi_{\mathrm{ch}}$ with (B) the minimum free energy path (MFEP) for stalk formation presented by Smirnova *et al.*,[1] computed along an order parameter given the three-dimensional (3D) hydrophobic membrane density and optimized with the string method. Curves in (B) were taken from Ref. 1. PMFs in panel (A) and MFEPs in (B) were computed with the same simulation systems and topologies, kindly provided by the authors of Ref. 1. (C) An older MARTINI POPC model with a 5-bead oleoyl tail was used, longer than the four-bead oleoyl model used for all other simulations of this study. The systems contained 128 POPC lipids per bilayer and either 200 (PChd200) or 220 (PChd220) water beads in the proximal compartment, corresponding to 1.56 or 1.72 water beads per lipid. Evidently, the PMFs along $\xi_{\mathrm{ch}}$ suggest lower free energies for the stalk as compared to the MFEPs along the density-based order parameter used in Ref. 1, possibly because the state with $\xi_{\mathrm{ch}} \approx 1$ includes far more conformational states than the well-defined 3D density specified by the final state $s \approx 1$ along string contour.[2]

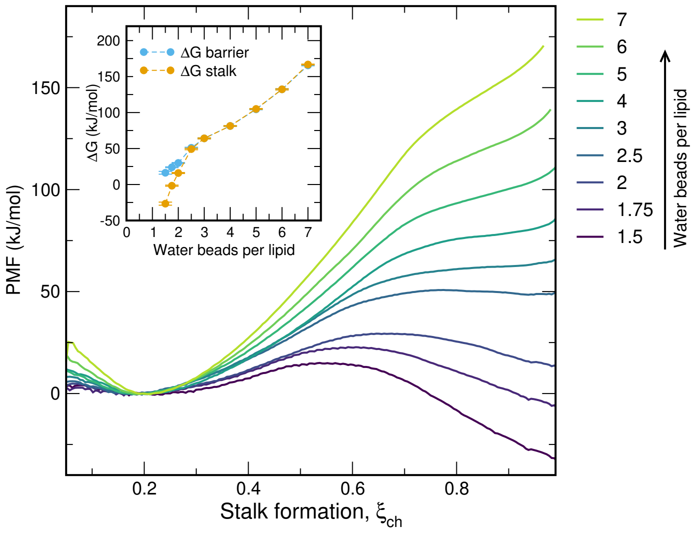


Figure S3: PMFs of stalk formation for membranes of pure POPC, computed with the beta-3.2 release of MARTINI 3.0. PMFs were computed with increasing amount of water in the proximal water compartment, defined by the number of water beads per lipid (see legend). Inset: Free energy of the stalk $\Delta G_{\mathrm{stalk}}$ and of the stalk nucleation barrier ($\Delta G_{\mathrm{barrier}}$, if present) versus water beads per lipid in the proximal compartment, as taken from the PMFs.

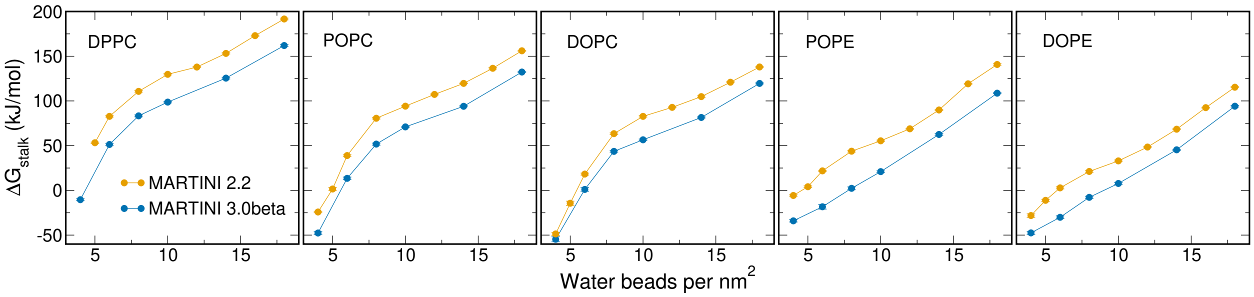


Figure S4: Free energies of stalk formation $\Delta G_{\mathrm{stalk}}$ for five lipid types (see labels) for various degrees of hydration in the proximal compartment. PMFs were computed with the Martini 2.2 model (yellow) or with the beta 3.2 release of Martini 3. The trends of $\Delta G_{\mathrm{stalk}}$ with hydration, tail unsaturation, and head group favorable agree among the two models. However, the beta release of Martini 3 yields systematically lower $\Delta G_{\mathrm{stalk}}$, implying more fusogenic membranes.

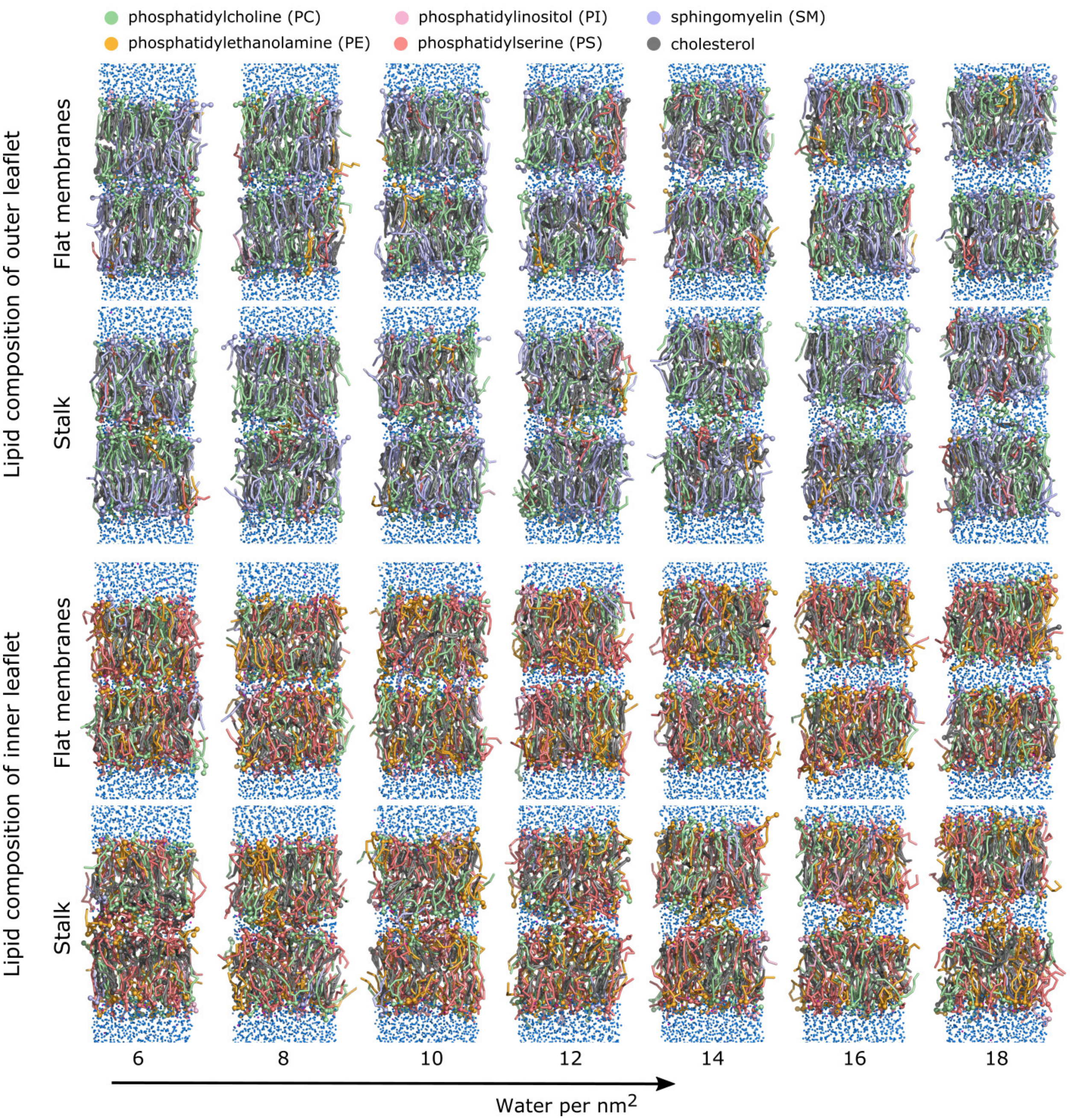


Figure S5: Simulations of stalk formation between membranes with the plasma membrane lipid composition. Upper two rows: lipid composition of the outer leaflet; lower two rows: lipid composition of the outer leaflet. From left to right: simulations systems with increasing hydration in the proximal water compartment, between 4 and 18 water beads per nm$^2$. Lipids are shown as sticks, water and $Na^+$ beads as blue and magenta spheres, respectively. The color of the lipids indicates the lipid type, see legend. Simulations frames were taken from the final snapshots of umbrella sampling simulations restrained to the state of two flat membranes ($\xi_{\mathrm{ch}} = 0.2$) or to the state of the open stalk ($\xi_{\mathrm{ch}} = 1$), respectively.

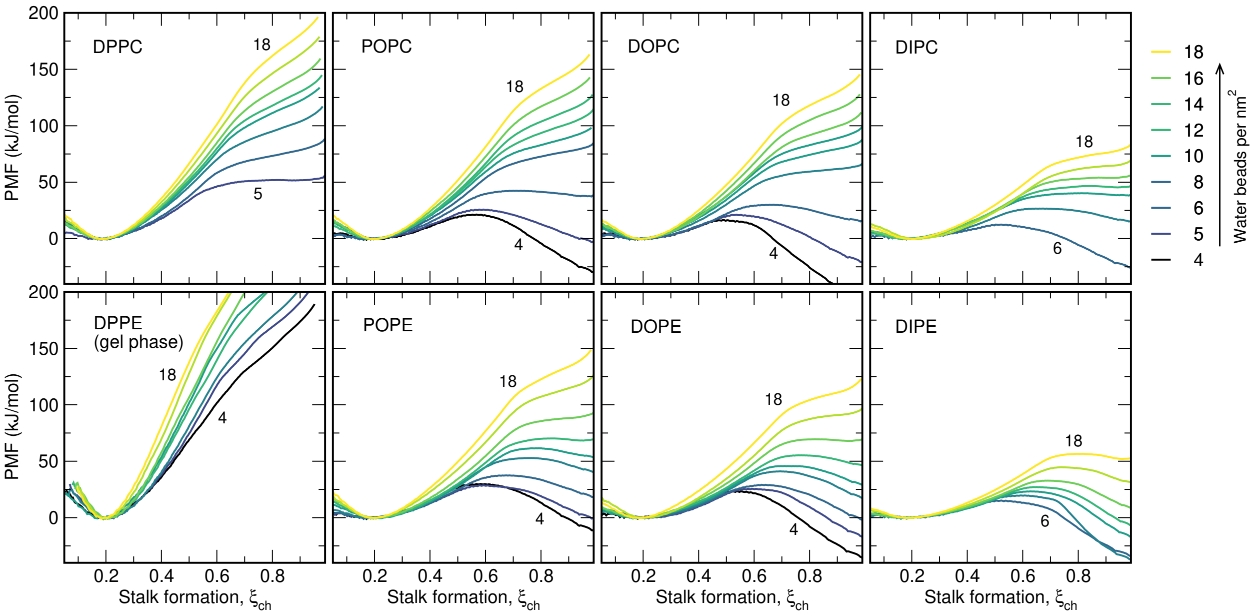


Figure S6: PMFs of stalk formation for PC lipids (upper row) and PE lipids (lower row) with increasing tail unsaturation (from left to right), computed with the MARTINI 2.2 model. $\Delta G_{\text{stalk}}$ for PE membranes is mostly lower as compared to PC membranes. Exception are the $\Delta G_{\text{stalk}}$ values at very low hydration, such as DOPC versus DOPE at 4–5 water beads/nm$^2$. Another exception is given by DPPE membranes, which formed a gel phase in the simulations, leading to greatly increased $\Delta G_{\text{stalk}}$ values and highly unstable stalks.

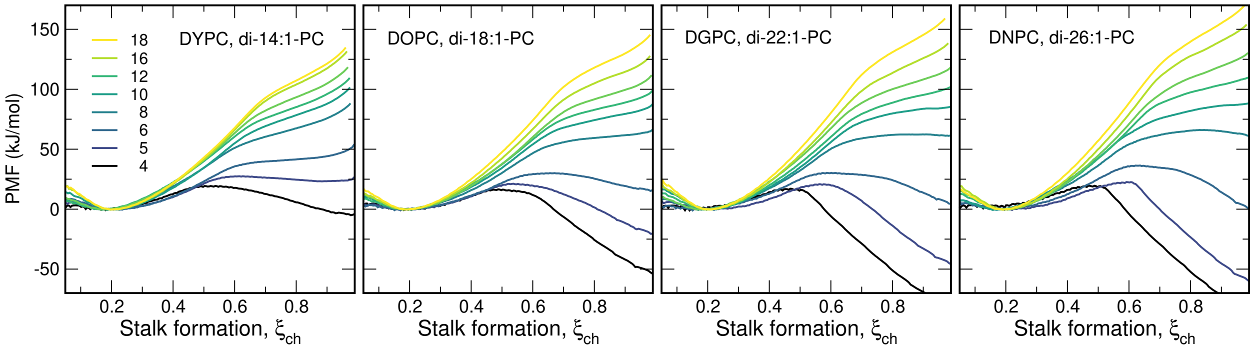


Figure S7: On the effect of tail length on the free energies of stalk formation: PMFs of stalk formation with membranes of (from left to right) DYPC, DOPC, DGPC, and DNPC. These MARTINI lipid types correspond approximately to atomistic lipids di-14:1-PC, di-18:1-PC, di-22:1-PC, and di-26:1-PC, respectively. The color code (from black to yellow) indicates increased hydration from 4–18 water beads per nm$^2$ in the proximal water compartment. Simulation frames of the open stalk for system with 6 water beads/nm$^2$ are shown in Fig. S9.

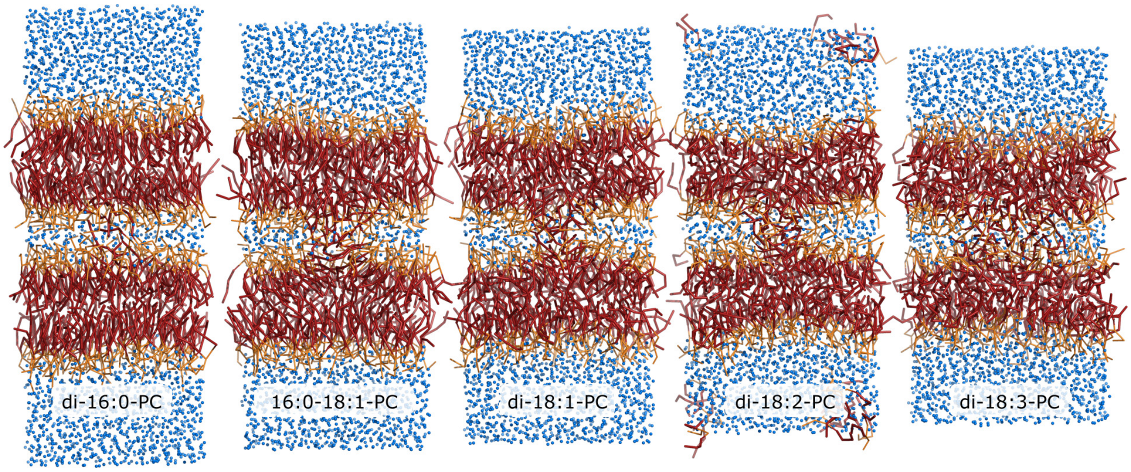


Figure S8: Typical simulation frame with an open stalk across membranes with increasing tail unsaturation. From left to right: of DPPC, POPC, DOPC, DLiPC, and DFPC. These MARTINI lipid types contain 4 beads and correspond approximately to atomistic lipids di-16:0-PC, di-16:0-18:1-PC, di-18:1-PC, di-18:2-PC, di-18:3-PC respectively. The proximal water compartment contains 6 water beads per $nm^2$. Frames were taken from the final snapshot of the last umbrella sampling window restrained to $\xi_{ch} = 1$. Lipid tails are shown as dark red sticks, head groups and glycerol region as orange sticks, and water beads as blue spheres.

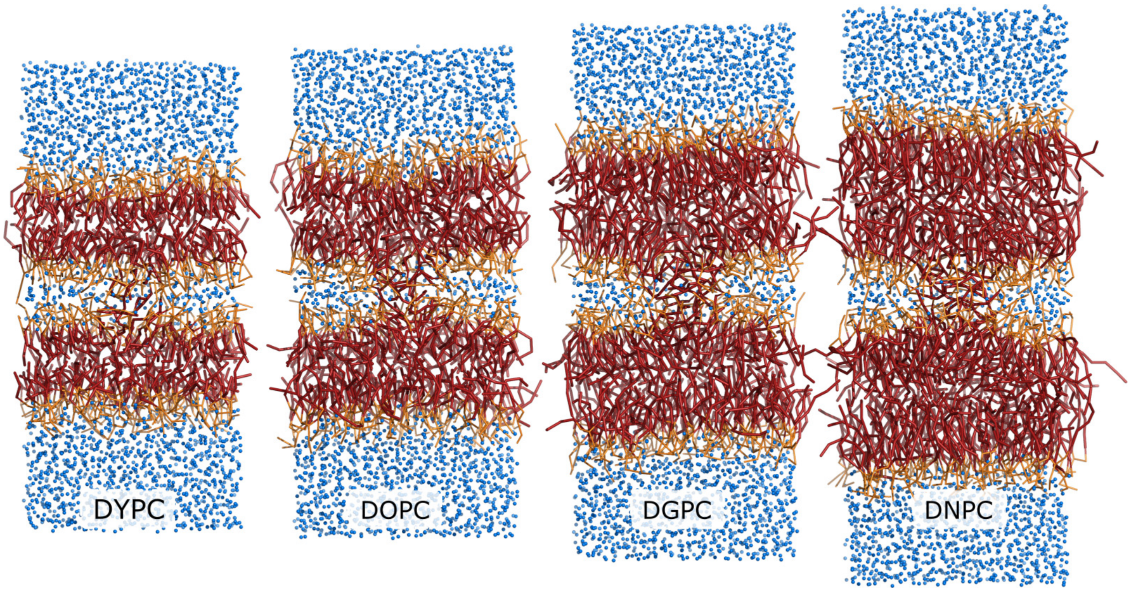


Figure S9: Typical simulation frame with an open stalk across membranes with increasing tail length. From left to right: of DYPC, DOPC, DGPC, and DNPC. These MARTINI lipid types correspond approximately to atomistic lipids di-14:1-PC, di-18:1-PC, di-22:1-PC, and di-26:1-PC, respectively. The proximal water compartment contains 6 water beads per $nm^2$. Frames were taken from the final snapshot of the last umbrella sampling window restrained to $\xi_{ch} = 1$. Lipid tails are shown as dark red sticks, head groups and glycerol region as orange sticks, and water beads as blue spheres.

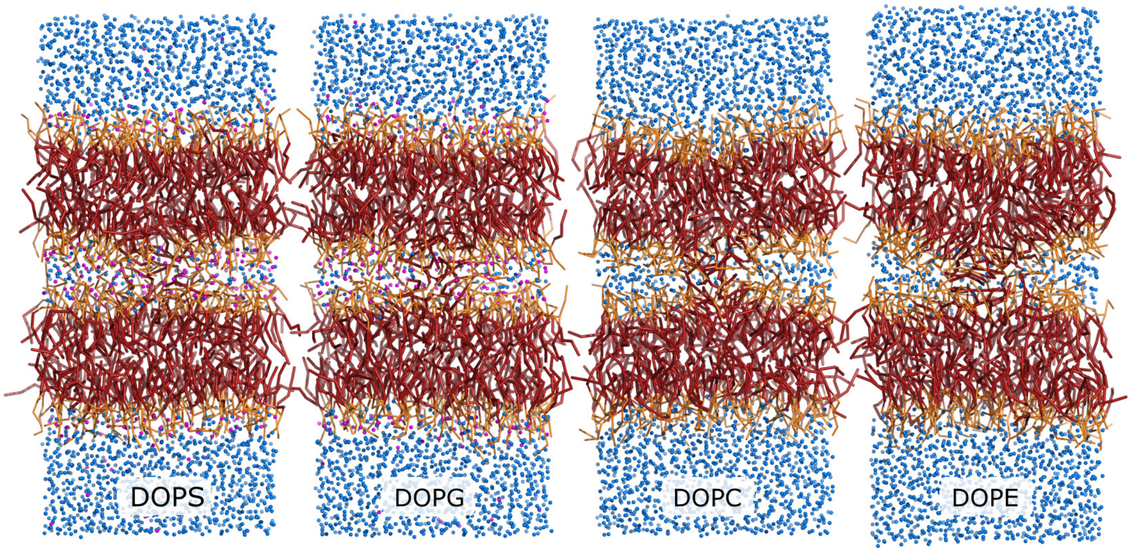


Figure S10: Typical simulation frame with an open stalk across membranes of (from left to right) DOPS, DOPG, DOPC, DOPE, taken from the final snapshot of the last umbrella sampling window restrained to $\xi_{ch} = 1$. Lipid tails are shown as dark red sticks, head groups and glycerol beads as orange sticks, water beads as blue spheres, and $Na^+$ beads as magenta spheres.

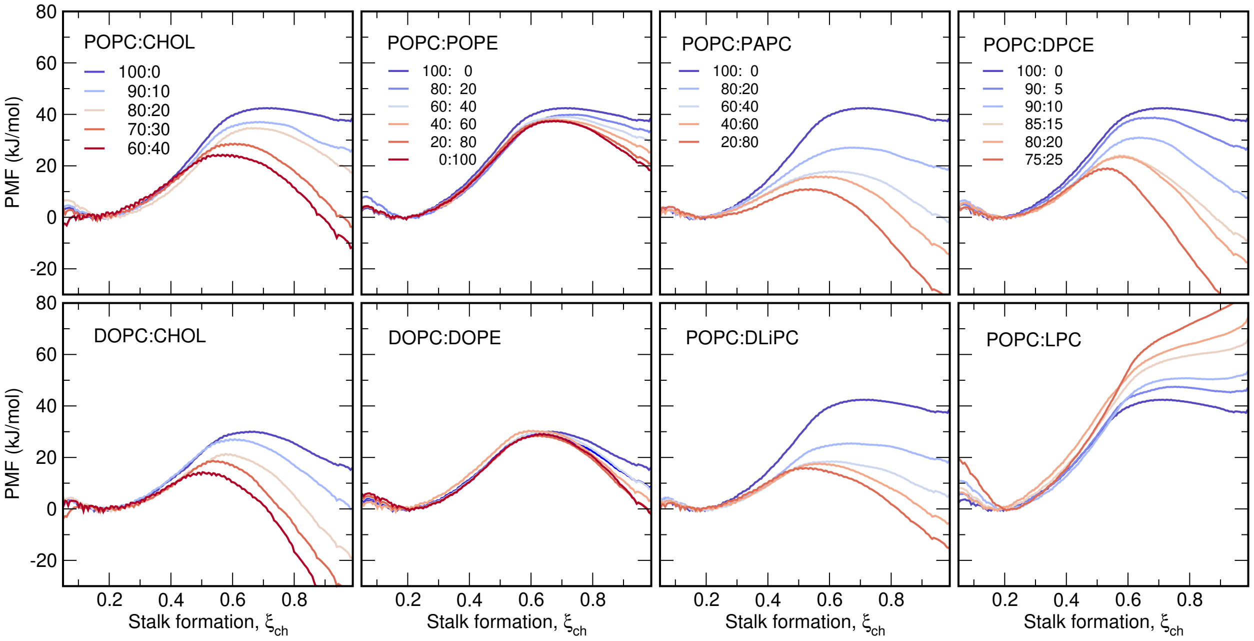


Figure S11: Typical PMFs of stalk formation for binary lipid mixtures, as indicated in the figure caption. Lipids are denoted with Martini lipid names. Lipid abbreviations are listed in the legend of Fig. 5.

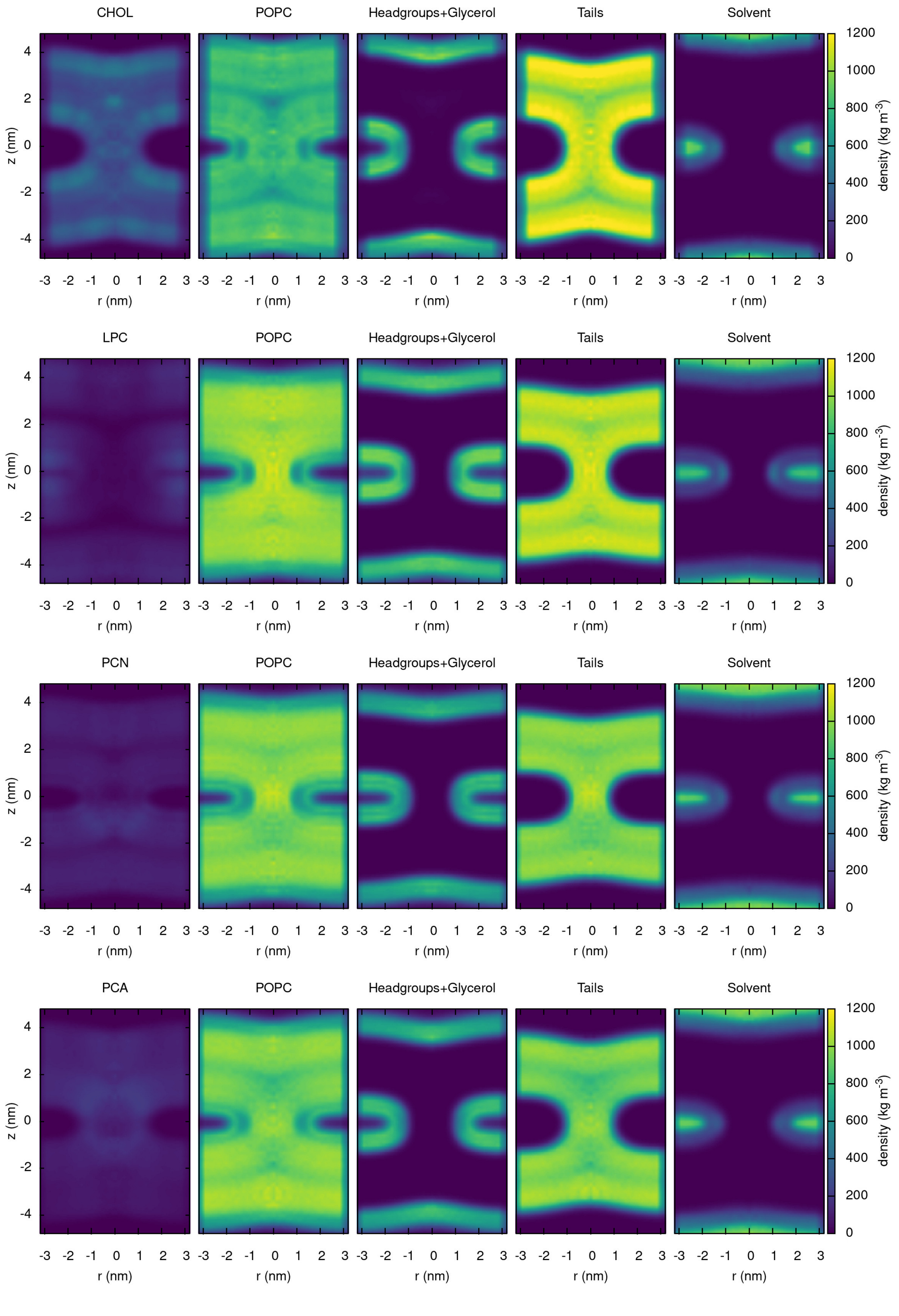


Figure S12: Mass densities at lipid stalk (set 1) for systems POPC:Cholesterol 80:20, POPC:LPC 80:20, POPC:PCN 80:20, POPC:PCA 80:20. Densities were computed from the umbrella window restrained to $\xi_{\mathrm{ch}} = 1$, omitting the first 50 ns for equilibration. The densities $\rho(r, z)$ were computed as function of the lateral distance $r$ and normal distance $z$ from the center of the stalk, defined as the center of the cylinder used to define $\xi_{\mathrm{ch}}$ (see Methods). $\rho(r, z)$ were copied to negative $r$-values purely for visualization purposes.

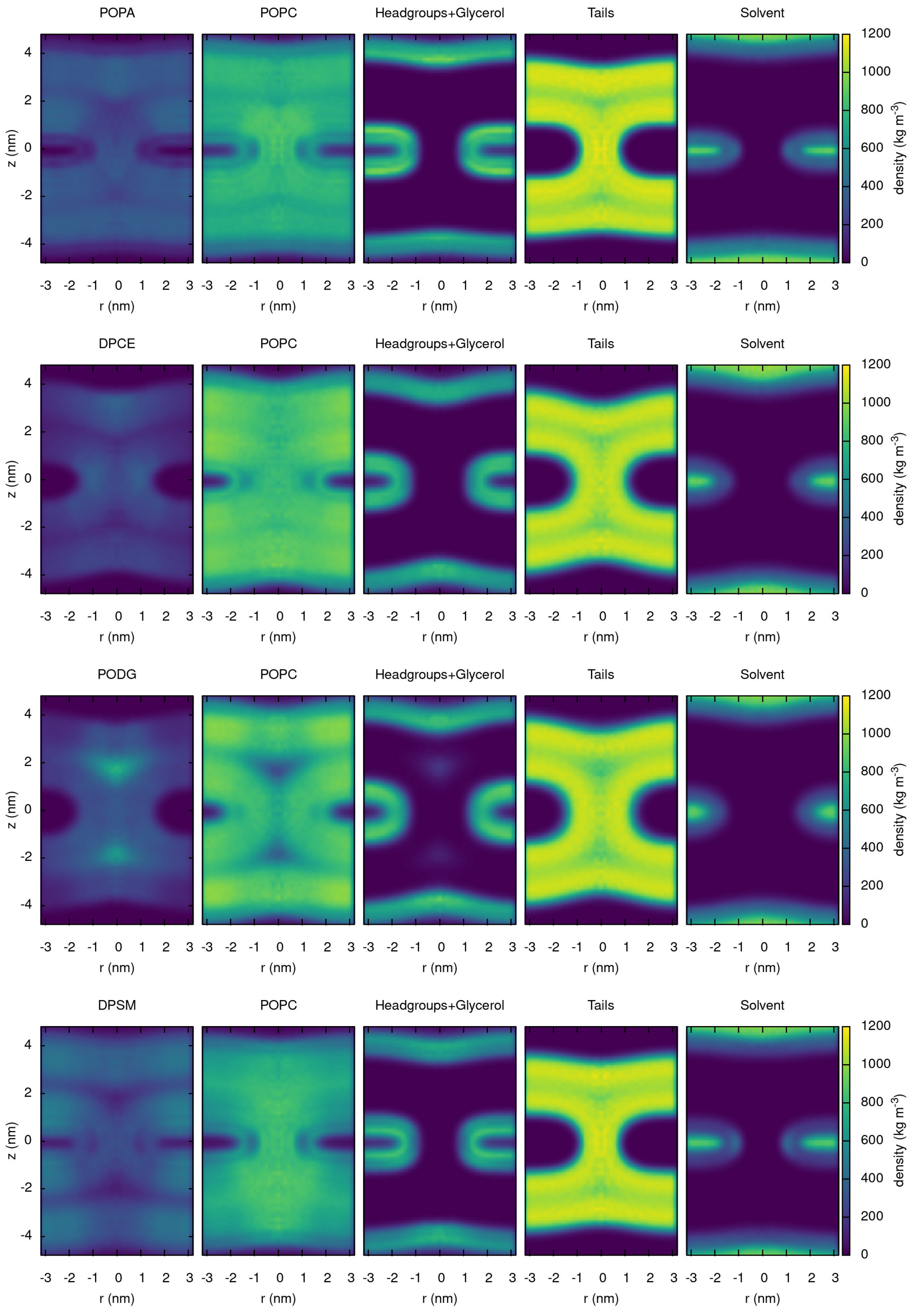


Figure S13: Mass densities at lipid stalk (set 2) for systems POPC:POPA 80:20, POPC:DPCE 80:20, POPC:PODG 80:20, POPC:DPSM 60:40.

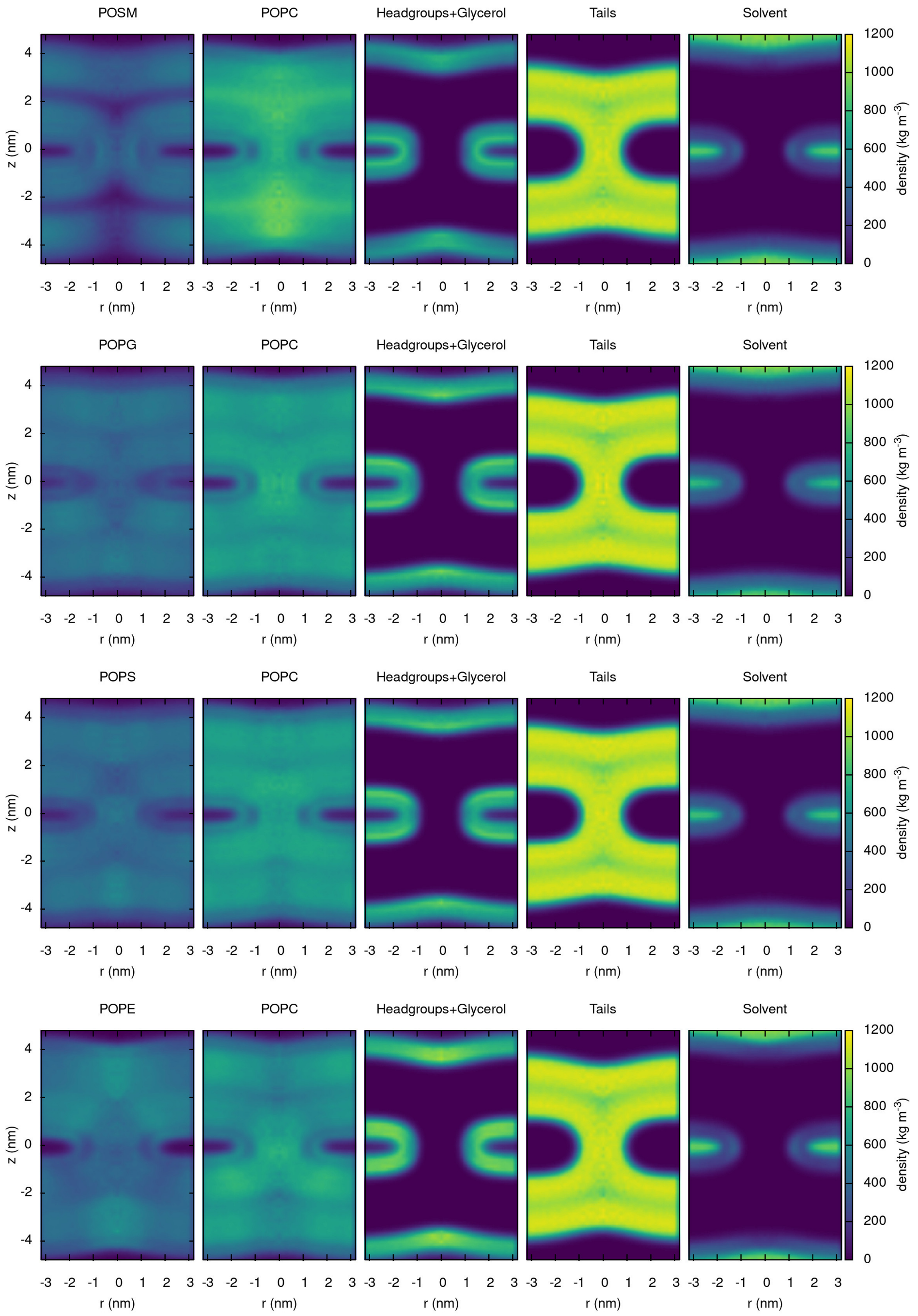


Figure S14: Mass densities at lipid stalk (set 3) for systems POPC:POSM 60:40, POPC:POPG 60:40, POPC:POPS 60:40, POPC:POPE 60:40.

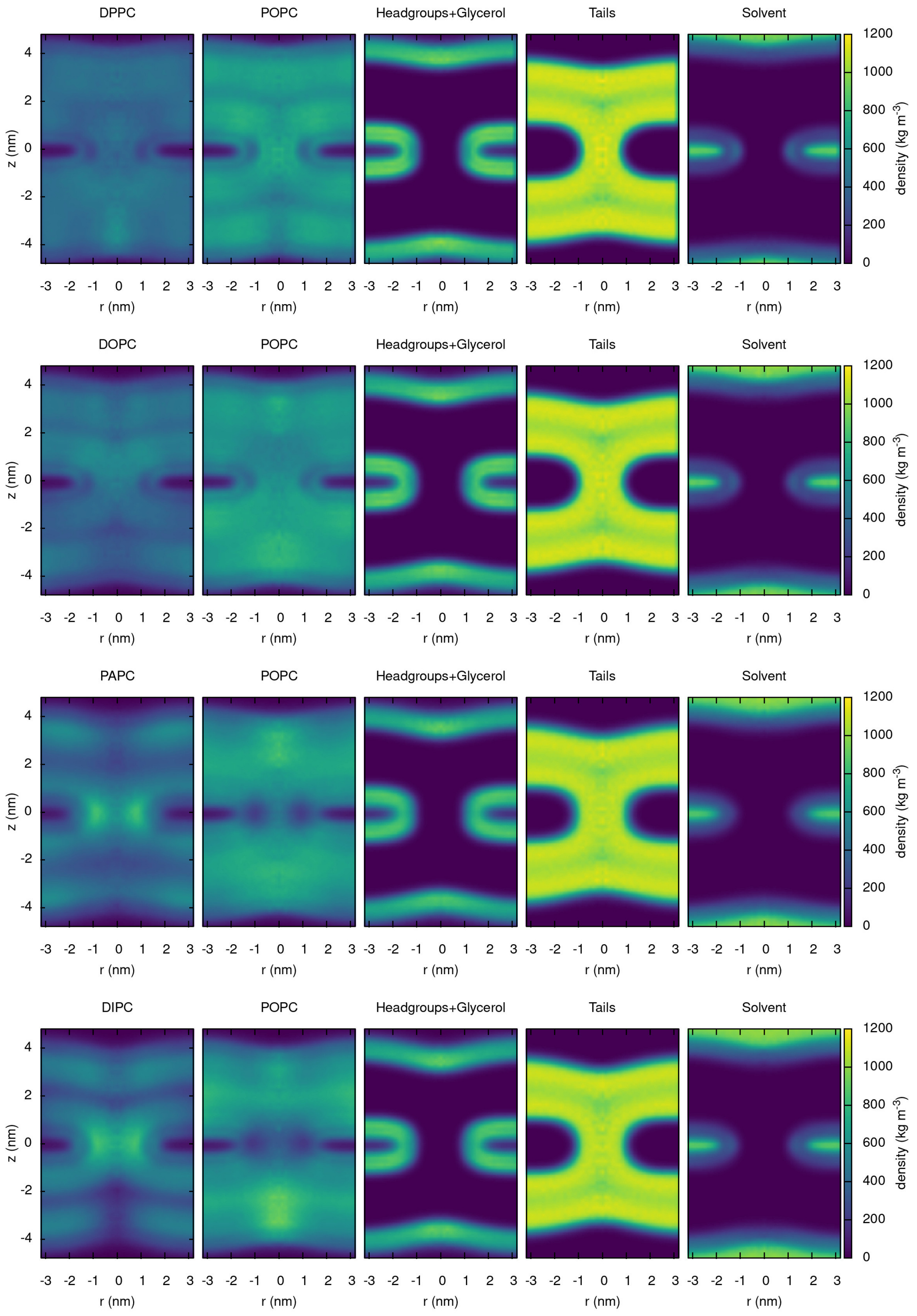


Figure S15: Mass densities at lipid stalk (set 4) for systems POPC:DPPC 60:40, POPC:DOPC 60:40, POPC:PAPC 60:40, POPC:DLiPC 60:40 (Martini name: DIPC).

Table S1: Lipid composition of plasma membrane models: MARTINI residue name, approximate atomistic correspondence, number of lipids per bilayer for the outer ($N_{\mathrm{outer}}$) and for the inner leaflet ($N_{\mathrm{inner}}$) models. Lipid types are: phosphatidylcholine (PC), sphingomyelin (SM), phosphatidylinositol (PI), phosphatidylethanolamine (PE), cholesterol (CHOL).

| MARTINI | ~atomistic | $N_{\mathrm{outer}}$ | $N_{\mathrm{inner}}$ |
|---|---|---|---|
| DPPC | di-16:0-PC | - | 2 |
| POPC | 16:0-18:1-PC | 6 | 4 |
| PIPC | 16:0-18:2-PC | 18 | 12 |
| PAPC | 16:0-20:4-PC | 6 | - |
| DPSM | di-16:0-SM | 14 | 2 |
| PNSM | 16:0-24:1-SM | 8 | - |
| PXSM | 16:0-24:0-SM | 8 | - |
| PAPI | 16:0-20:4-PI | 2 | 2 |
| PAPS | 16:0-20:4-PS | 4 | 30 |
| POPE | 16:0-18:1-PE | - | 2 |
| PAPE | 16:0-20:4-PE | 2 | - |
| PUPE | 16:0-22:6-PE | - | 12 |
| OAPE | 18:1-20:4-PE | - | 4 |
| CHOL | | 52 | 52 |
| Total | | 120 | 122 |

### Supporting Methods

#### Simulation setup and parameters

The double-membrane systems were set up with a multi-step protocol that ensures the requested degree of hydration between the two membranes, irrespective of the lipid composition. First, a single membrane system was set up with the Insane software.[3] The membrane was hydrated with the requested number of water beads and neutralized with sodium beads, as needed. The energy of the system was minimized and the membrane was equilibrated for 20 ns to fully relax the box dimensions.

Next, two copies of the membrane were stacked on top of each other, (i) leading to the requested degree of hydration between the proximal leaflets of the double-membrane and (ii) to balanced electric charges between the two water compartments, thereby avoiding a transmembrane electric potential. To fully hydrate the distal leaflets, the box was enlarged along the $z$ direction and additional water was added to the distal water compartment, until the distal leaflets were hydrated with 10 water beads per lipid. The double membrane was equilibrated for another 20 ns.

In long simulations of the double-membrane system, we observed occasional membrane permeation by water beads, which would change the degree of hydration of the proximal leaflets. To avoid such permeation, we applied flat-bottomed position restraints (FB-posres) to all water beads. The reference positions of all FB-posres were taken as the center of the box along $z$. For water in the proximal compartment, an attractive FB-posres was applied, for which the thickness of flat region was set to $z_{\mathrm{fb}} = (z_u - z_l)/2 - 0.5\,\mathrm{nm}$, where $z_u$ and $z_l$ denote the center of mass positions of the upper and lower membrane, respectively. For water in the distal compartment, a repulsive FB-posres was applied, where the repulsive region had a thickness of $(z_u - z_l)/2 + 0.5\,\mathrm{nm}$. The force constant for the quadratic potential was set to $100\,\mathrm{kJ\ mol^{-1}nm^{-2}}$. Together, these FB-posres potentials allowed normal diffusion and fluctuation of water in both compartments and applied only if water beads deeply penetrated into the hydrophobic membrane cores.

Unbiased simulations were carried out with the Gromacs simulation software, version 2020.3.[4] If not stated otherwise, interaction potentials were described with the MARTINI model 2.2.[5] To test the influence of the MARTINI model generation, simulations for certain lipids were also carried out with the beta release 3.2 of Martini 3.0. Neighbor lists were updated with the Verlet algorithms. Lennard-Jones and Coulomb potentials were truncated at 1.1 nm. The temperature was controlled at 310 K (if not stated otherwise) through velocity rescaling using four separate coupling groups for the two membranes and for the two water compartments ($\tau = 0.1$ ps).[6] The pressure was kept at 1 bar with the Berendsen barostat ($\tau = 6$ ps).[7] The integration time step was set to 20 fs or 30 fs for simulations with or without cholesterol, respectively.

#### Reaction coordinate for stalk formation

PMFs were computed along the "chain coordinate" $\xi_{\text{ch}}$, which was originally introduced to drive pore formation in membranes.[8,9] $\xi_{\text{ch}}$ quantifies the connectivity of two compartments of specific atoms. $\xi_{\text{ch}}$ was defined by a cylinder of radius 1.2 nm that spans the two hydrophobic regions of the two membranes and the proximal water compartment (Figure S1). The cylinder is decomposed into slices with a thickness of 1Å. Then, $\xi_{\text{ch}}$ is approximately given by the fraction of slices that are filled by lipid tail beads:

$$\xi_{\text{ch}} = \frac{1}{N_s} \sum_{s=0}^{N_s-1} \delta_\zeta(n_s^{(t)}) \tag{1}$$

Here, $N_s$ is the number of cylinder slices, $n_s^{(t)}$ the number of tail beads in slice $s$, and $\delta_\zeta$ is a (differentiable) indicator function that takes $\delta_\zeta = 0$ for an empty slice ($n_s^{(t)} = 0$) and $\delta_\zeta \approx 1$ for a filled slice ($n_s^{(t)} \geq 0$). Hence, upon pulling the system along $\xi_{\text{ch}}$, the slices are filled one-by-one, thereby gradually forming a hydrophobic connection between the two hydrophobic membrane cores, as required for stalk formation. As bead contributing to $\xi_{\text{ch}}$, hydrophobic lipid tail beads as well as hydrophobic beads of cholesterol were used.

To render $\xi_{\mathrm{ch}}$ differentiable, the function $\delta_\zeta$ was defined with a differentiable switch function that approximates and indicator function:[8]

$$\delta_\zeta(x) = \begin{cases} \zeta x & \text{if } x \leq 1 \\ 1 - c\,\mathrm{e}^{-bx} & \text{if } x > 1 \end{cases} \tag{2}$$

Here, the parameter $\zeta$ indicates the fraction to which the slice is filled upon the addition of the first apolar bead into the slice. We used $\zeta = 0.75$ in this study. The parameters $b$ and $c$ are taken as $b = \zeta/(1-\zeta)$ and $c = (1-\zeta)\mathrm{e}^b$, leading to a continuous and differentiable switch function. Likewise, the number of beads $n_s^{(t)}$ in slice $s$ was defined with a differentiable indicator function:

$$n_s^{(t)} = \sum_{j=1}^{N_b} f(\mathbf{r}_j) \tag{3}$$

Here, the sum is taken over all $N_b$ apolar beads and $\mathbf{r}_j$ denotes the Cartesian coordinates of bead $j$. $f$ is a three-dimensional indicator function that takes unity inside the volume of slice $s$, and $f$ smoothly switches to zero at the slice boundaries.

Critically, the lateral position of the cylinder was not fixed but dynamically defined to allow the cylinder to "follow" the stalk as the stalk travels parallel to the membrane plane. This property excludes that the system moves along $\xi_{\mathrm{ch}}$ by shifting the stalk laterally out of the cylinder, which would lead to undesired hysteresis effects.[10] For details, we refer to previous work.[8] To visualize the stalk in molecular graphics, the stalk was translated to the box center purely for illustration purposes (Figs. S1, S5, S9, S10).

The number of slices $N_s$ and thereby the height of the cylinder was chosen depending on the thickness of the proximal water compartment. To this end, the average $\xi_{\mathrm{ch}}$ was computed from at last 10 ns of an equilibrium simulation of the flat double membrane using various $N_s$ between 8 and 45, and employing the 'rerun' functionality of the Gromacs mdrun module. Henceforth, $N_s$ was chosen such that $\xi_{\mathrm{ch}} \approx 0.2$ for the flat membrane. In other words, $N_s$ was chosen such that ∼20% of the slices were filled by lipid tail beads in a flat membrane. For membranes of pure POPC and 4 to 18 water beads per $nm^2$ in the proximal compartment, for instance, this protocol led to cylinders with 16 to 36 slices.

#### Umbrella sampling simulations

PMFs were computed using umbrella sampling (US). Initial frames for US were taken from constant-velocity pulling simulations, in which the systems were pulled from $\xi_{ch} = 0.1$ to $\xi_{ch} = 1$ within 200 ns, using a force constant of 3000 kJ/mol. Visual inspection of the simulations showed that pulling $\xi_{ch}$ led to gradual stalk formation in all membrane system. 19 umbrella windows were used with reference positions between 0.1 and 1 in steps of 0.05. The force constant was set to 3000 kJ/mol. Each window was simulated for 200 ns, where the first 50 ns were omitted for equilibration. An integration time step of 20 fs was used. All other parameters were chosen as described above. The PMFs were computed with the weighted histogram analysis method (WHAM),[11] as implemented in the gmx wham module of Gromacs.[12] Statistical errors were estimated with the Bayesian bootstrap of complete histograms. Accordingly, in each round of bootstrapping, random weights were assigned to all histograms, and the randomly weighted histograms were used to compute a bootstrapped 'synthetic' PMF. Each bootstrapped PMF was defined to zero at $\xi_{ch} = 0.2$ before computing the standard deviation among the bootstrapped PMFs. The procedure suggested statistical errors in the order of 1 to 3 kJ/mol, indicative of well converged PMFs.

The free energy of stalk formation $\Delta G_{stalk}$ was defined as the average of the PMF in the interval $\xi_{ch} \in [0.95, 1]$.

#### Absence of hysteresis between stalk opening and closing pathways

As a test for the validity of the reaction coordinate, we computed the PMFs along stalk-opening and stalk-closing pathways (Fig. S16). We obtained nearly identical PMFs along opening and closing pathways, confirming the absence of undesired hysteresis.

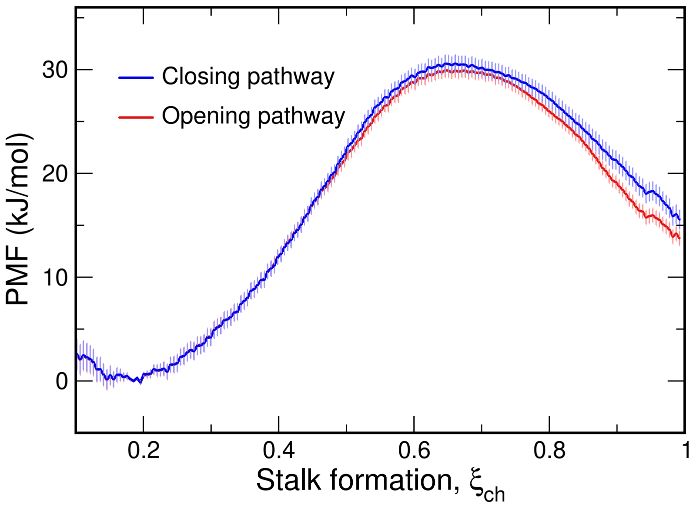


Figure S16: PMFs of stalk formation for membranes of pure POPC with 2 water beads per lipid in the proximal compartment. Starting frames for umbrella sampling were taken from constant-velocity pulling simulation conducted either in stalk-opening direction (red) or in stalk-closing direction (blue). The absence of any hysteresis suggests that the PMFs are converged and that the pathways are reversible.

#### Unbiased simulations of stalk formation and closure

To test whether the PMF along $\xi_{\mathrm{ch}}$ reflects the true free energy difference between the flat membrane and the stalk, and to obtain the rates of stalk formation and closure, we carried out unbiased simulations. Here, we used the beta-3.2 release of MARTINI 3.0. We simulated a double-membrane of pure POPC with 230 and 1920 water beads in the proximal and distal water compartments, respectively, for which the PMF suggested a free energy difference between stalk and flat membrane of $\Delta G_{\mathrm{stalk}} \approx 0$ (Fig. 2). Four replicas of 200 µs each were simulated, which carried out 8 transitions of stalk formation and 7 transitions of stalk closure corresponding to rates of $k_{\mathrm{stalk}} = 16\,\mathrm{ms}^{-1}$ and $k_{\mathrm{closure}} = 23\,\mathrm{ms}^{-1}$, respectively. Hence, the free simulations suggest a free energy of stalk formation of $\Delta G_{\mathrm{stalk}} = -k_B T \ln(k_{\mathrm{stalk}}/k_{\mathrm{closure}}) = 0.9\,\mathrm{kJ/mol}$, in excellent agreement with the PMF.

#### Attempt frequency of stalk formation

According to transition state theory, the rate of barrier crossing is given by

$$k = \nu \, \mathrm{e}^{-\Delta G^{\ddagger}/k_B T}, \tag{4}$$

where $\nu$ is the attempt frequency and $\Delta G^{\ddagger} = 25\,\mathrm{kJ/mol}$ the barrier height in the PMF (Fig. 2B). With the observed rate of approximately $20\,\mathrm{ms}^{-1}$ and the applied temperature of 310 K, this suggests an attempt frequency of $\nu \approx 0.3\,\mathrm{ns}^{-1}$ or, equivalently, approximately one attempt per 3 ns.

#### Density calculations

The mass density around the stalk was computed with an in-house modification of the Gromacs module gmx density. For the density calculations only, the masses of cholesterol beads were modified to resemble the physical mass distribution, taking the common mapping of three to four heavy atoms onto one CG bead. This step was necessary because the masses of the original MARTINI cholesterol model have been optimized to reproduce the moments of inertia, but not the physical mass distribution. For all other lipids, the original MARTINI mass beads were used.
